# Elevation of the mechanically-sensitive protein emerin links nuclear mechanotransduction to tau-induced cytoskeletal remodeling in neurons

**DOI:** 10.1101/2025.10.15.682650

**Authors:** Claira Sohn, Sammy Pardo, Dana Molleur, Satvik R Paduri, Morgan Lambert, Morgan G Thomas, Erich J Sohn, Susan T Weintraub, Bess Frost

## Abstract

Tauopathies are a group of neurodegenerative disorders, including Alzheimer’s disease, that are neuropathologically defined by deposition of pathological forms of tau in the brain. While tau is reported to drive neurotoxicity by negatively affecting cytoskeletal, nucleoskeletal, and genomic architecture, the mechanisms mediating tau-induced dysfunction of the cytoskeleton and nucleoskeleton are incompletely understood. Based on proteomic profiling, we identify a suite of cytoskeletal and nucleoskeletal proteins with differing abundance in a cellular model of tauopathy, iTau. Building upon previous findings that pathogenic forms of tau reduce nuclear tension, we find that protein levels of emerin, a central regulator of nuclear mechanotransduction, are significantly elevated in iTau cells and in induced pluripotent stem cell (iPSC)-derived neurons carrying a mutation in the microtubule-associated protein tau (*MAPT*) gene that causes autosomal dominant frontotemporal dementia. We find that neuronal emerin overexpression is sufficient to drive neurotoxicity, increase overall levels of filamentous actin (F-actin), and induce nuclear invagination, cellular phenotypes that also occur in settings of tauopathy. Mass spectrometry-based identification of emerin-interacting proteins in iTau-derived neurons reveals increased interactions with cytoskeletal proteins and reduced interactions with nuclear proteins. Indeed, we find that emerin relocalizes from the nucleus to the cytosol in the setting of tauopathy, suggesting that pathogenic tau impacts nuclear mechanotransduction pathways. Overall, we identify emerin as a mediator of cytoskeletal remodeling in tauopathy and provide a foundation for future studies into the mechanosensitive function of emerin in neurons.

**SIGNIFICANCE STATEMENT:** Cells experience and respond to diverse mechanical forces that shape their morphology, function, and survival through a process termed “mechanotransduction.” While well studied in non-neuronal cells, neuronal mechanotransduction remains poorly understood despite exposure of the brain to vascular flow, movement, injury, and disease. We identify the mechanosensitive protein emerin as a key regulator of nuclear mechanotransduction in neurons. Emerin overexpression is sufficient to increase filamentous actin, induce nuclear invagination, and drive neurotoxicity, revealing a novel function for emerin in neurons. In cellular models of tauopathy, emerin is elevated and relocalizes from the nucleus to the cytoplasm, where it alters cytoskeletal structure. These findings establish emerin as a mechanosensitive regulator in neurons and link disrupted nuclear neuronal mechanotransduction to neurodegeneration.

## INTRODUCTION

Cells of living organisms are subject to a variety of mechanical forces that influence cellular differentiation, size, morphology, and proliferation through a process termed mechanotransduction [1–3]. While mechanotransduction pathways have been primarily studied in non-neuronal cells, the brain is subject to acute and chronic changes in mechanical force due to blood flow through the vasculature, head injuries, aging, neurodegenerative processes, and other factors [4–6]. Studies in cultured rat cortical neurons suggest that neuronal networks have the capacity to adjust electrophysiological activity and cytoskeletal structure in response to mechanical force [7].

The nucleus is a central regulator of mechanotransduction [8–12]. Forces in the cytosol are transmitted to the nucleus through the nuclear membrane-spanning linker of nucleoskeleton and cytoskeleton (LINC) complex [11]. The LINC complex binds to actin and microtubules on the external face of the nuclear envelope and to lamin proteins along the internal face of the nuclear envelope. Lamin proteins assemble to form a meshwork of intermediate filaments that provide strength and structure to the nucleus. Lamina-associated polypeptide 2 (LAP2), emerin, and inner nuclear membrane protein Man1 (MAN1) (LEM)-domain proteins bind to the lamin nucleoskeleton, where they sense and respond to forces on the nucleus [8,13,14].

A growing body of work suggests that nuclear form and function are disrupted in the context of various neurodegenerative disorders, including Alzheimer’s disease and related tauopathies. Mechanistically, pathological forms of tau are reported to drive nuclear invagination due to their effects on the actin cytoskeleton [15] and microtubule network [16]. Studies in *Drosophila* suggest that tau-induced increases in F-actin [17] cause clustering of the LINC complex, which drives reduction of lamin protein levels and consequent nuclear pleomorphism [15]. Previous studies report that brains of patients with Alzheimer’s disease have reduced stiffness compared to age-matched controls, with regional stiffness further changing as the disease advances [18,19]. Based on these studies, alongside our previous finding that cultured cells harboring pathogenic tau have an overall decrease in nuclear tension [20], we hypothesized that pathways regulating nuclear mechanotransduction may be disrupted in tauopathy.

We find that pathological tau significantly alters total abundance of proteins involved in cytoskeletal and nuclear function, including the mechanically-sensitive protein emerin. Studies in fibroblasts indicate that emerin is necessary for proper cytoskeletal and nuclear mechanics [21–23]. While there are few studies investigating the role of emerin in post-mitotic cells, recent findings suggest that emerin is elevated in neurons in response to sensory input and regulates activity-induced neuronal plasticity in mice [24]. We find that emerin overexpression in cultured neurons is sufficient to increase F-actin, drive nuclear invagination, and induce neurotoxicity. In the setting of tauopathy, we find that emerin relocalizes from the nucleus to the cytoplasm, where it has increased interaction with cytoskeletal proteins and alters F-actin structure. Our findings lay the groundwork for future studies on the role of emerin and altered nuclear mechanotransduction in neurodegenerative tauopathies and highlight an emerging function of emerin as a regulator of nuclear mechanotransduction in neurons.

## RESULTS

### Proteomic profiling reveals cytoskeletal and nucleoskeletal regulators affected by pathogenic tau

The BE(2)-C iTau cellular model recapitulates the negative effects of tau on nuclear morphology [20] described in other laboratory models of tauopathy and in human disease [15,16,25–27]. iTau cells feature doxycycline-inducible expression of human tau carrying the frontotemporal dementia-associated [28] *MAPT* mutation R406W (tau^R406W^). We have previously used iTau cells to discover that pathogenic forms of tau decrease nuclear tension [20]. The inducible green fluorescent protein (iGFP) model, which does not accumulate disease-associated tau phosphoepitopes, serves as a control for transgenic protein overexpression. We leveraged iTau and iGFP cells in an unbiased approach to discover potential novel mechanisms underlying tau-induced cytoskeletal and nuclear dysfunction using data-independent acquisition (DIA) high-performance liquid chromatography (HPLC) mass spectrometry.

Proteomic analysis detects approximately 5,300 proteins per sample, with high group correlation among iGFP and iTau replicates (**Supplemental Data 1**). We identify 290 significantly differentially abundant proteins in iTau vs. iGFP cells with a fold change of 1.5 or greater (**Fig. 1A**). Pathway analysis reveals a significant increase of proteins involved in metabolic pathways in iTau cells (**Fig. 1B**), consistent with prior reports of disrupted lipid metabolism and mitochondrial function in tauopathy [29–33]. Proteins involved in pathways regulating synaptic interactions, membrane trafficking, and vesicle transport are depleted in iTau cells, also consistent with previous work [34–36].

**Figure 1:**
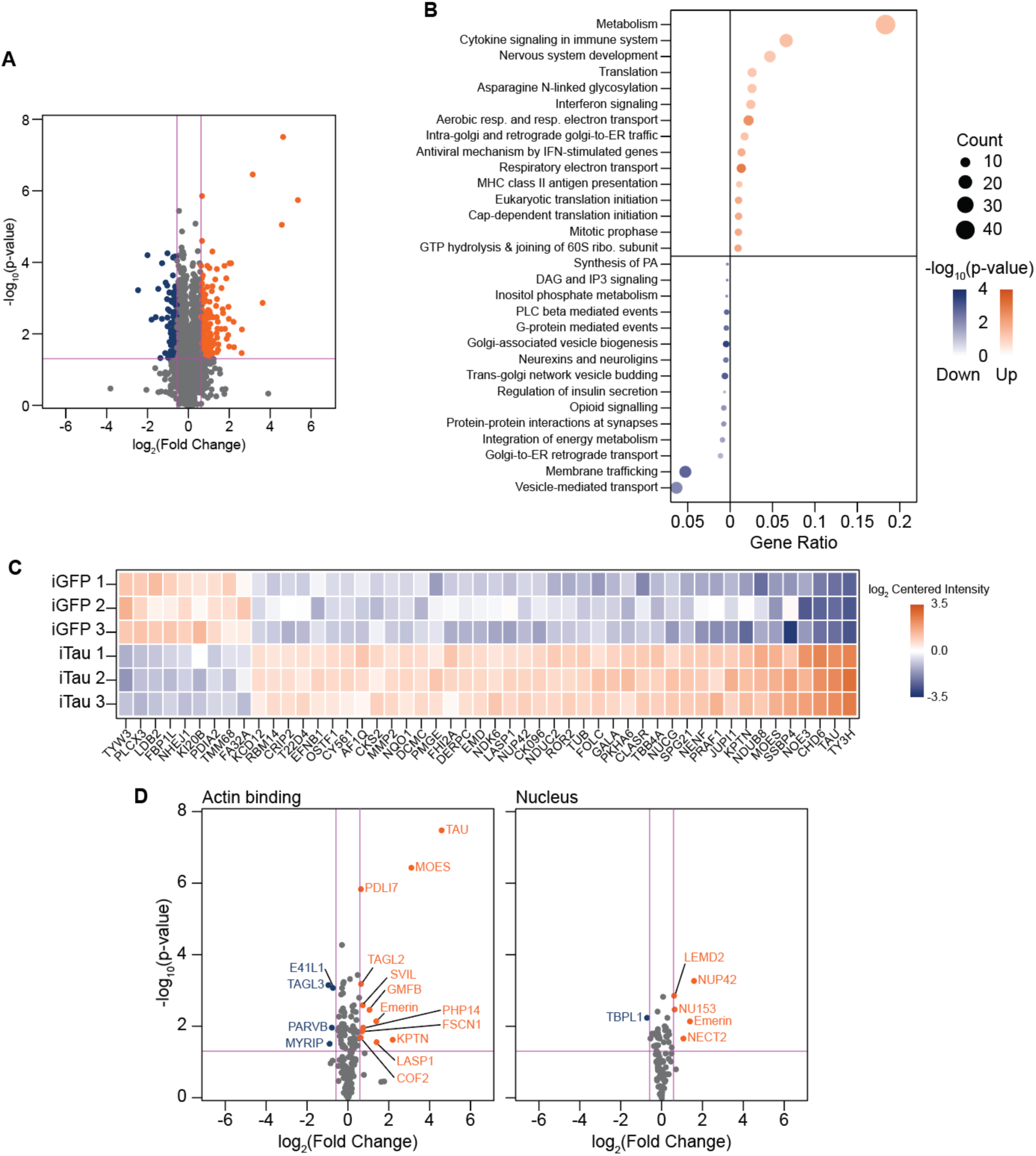
Proteomic profiling of iGFP and iTau cells reveals cytoskeletal and nucleoskeletal regulators affected by pathogenic tau. (**A**) Volcano plot showing differentially expressed proteins in iTau cells compared to iGFP based on mass spectrometry. (**B**) Pathway analysis of significantly altered pathways in iTau cells. (**C**) Top 50 significant proteins with the largest change in iTau cells. (**D**) Volcano plots of actin-binding (GO:0003779) or nuclear proteins (GO:0005634) in iTau cells compared to iGFP. n=3 biological replicates per group. Log_2_FC for significance is +/- 0.585 and p≤0.05.

Relevant to our focus on the effects of tau on the cytoskeleton and nucleoskeleton, we find many proteins involved in cytoskeletal and nuclear function among the top 50 differentially expressed proteins in iTau cells (**Fig. 1C**). Focusing on gene ontology (GO)-defined “actin-binding” and “nucleus” terms, we find that differentially expressed actin-binding proteins and nuclear proteins generally skew towards increased expression in iTau cells; differentially expressed nuclear proteins relate to nuclear trafficking, nuclear mechanotransduction, and nuclear morphology (**Fig. 1D**).

Emerin is among the most significantly elevated actin-binding and nuclear proteins in iTau cells. Studies in non-neuronal cells identify emerin as a mechanically sensitive protein that responds to changes in cytoskeletal force and nuclear tension by shifting its localization from the nucleus to the cytosol [8,37–39], and regulates mechanotransduction through interactions with the nucleo- and cytoskeleton [23,21]. While tau-induced elevation of emerin is highly relevant to our previous discovery that pathogenic forms of tau alter nuclear tension [20], little is known regarding the role of emerin as a mechanotransducer specifically in neurons. Given its significant upregulation in iTau cells, mechanosensitive properties, and dual role at the nucleo/cytoskeletal interface, we focused on emerin as a potential mediator of tau-induced changes in the cytoskeleton and nucleoskeleton.

### Emerin protein levels are elevated in neurons harboring pathological tau

As the BE(2)-C neuroblastoma cell line used to generate iTau and iGFP cells retains proliferative capacity and lacks key features of mature neurons, we induced cell cycle exit and neuronal differentiation by treating cells with 10 μM retinoic acid for seven days. Emerin protein is significantly elevated in iTau neurons based on immunofluorescence (**Fig. 2A**) and Western blotting (**Fig. 2B**).

**Figure 2:**
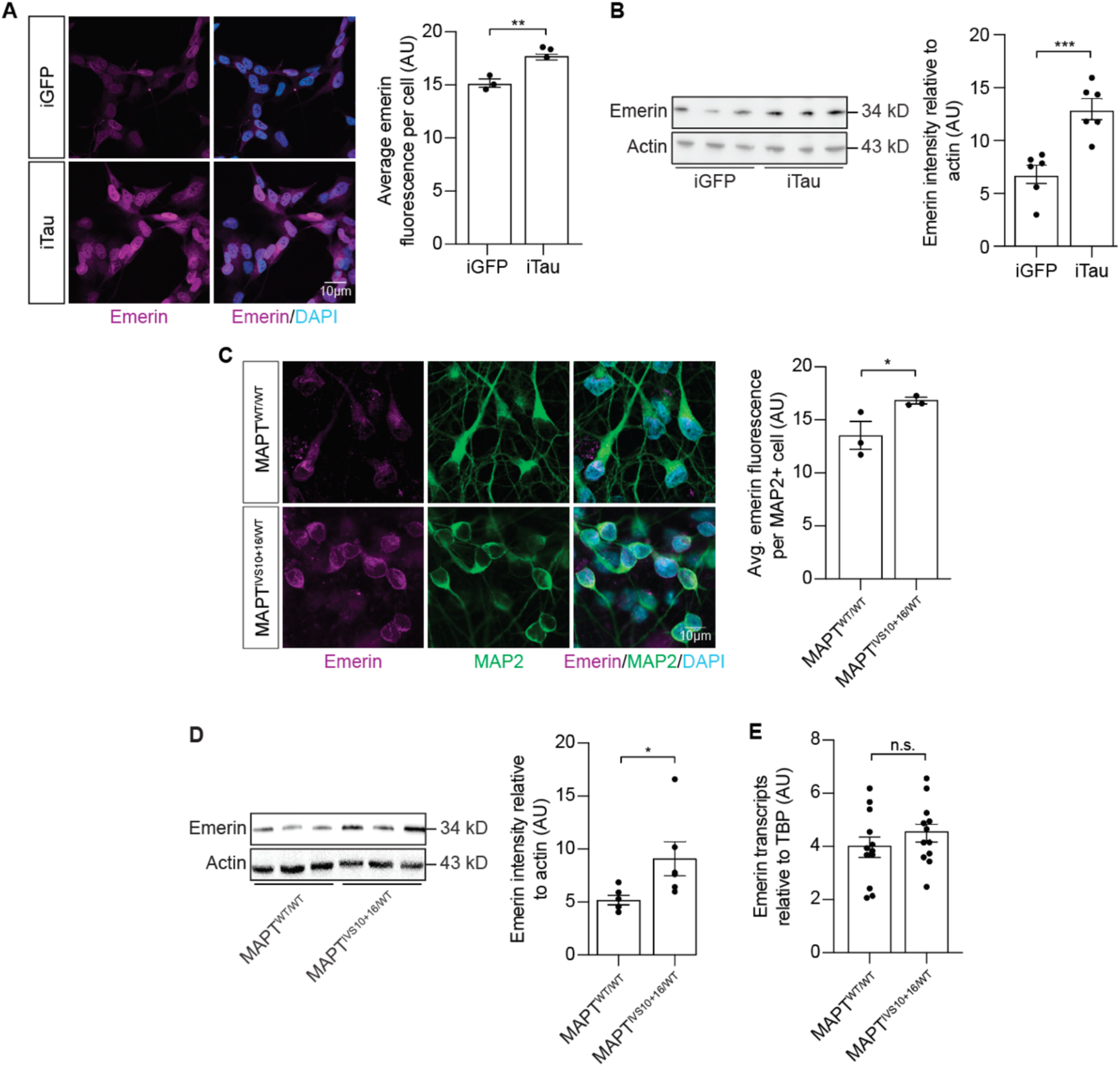
Emerin protein levels are elevated in neurons harboring pathological tau. Significant increase in emerin in iTau neurons based on (**A**) immunofluorescence and (**B**) Western blotting after one week of retinoic acid-mediated neuronal differentiation and 24-hour doxycycline-mediated induction of tau or GFP expression. Significant increase in emerin in *MAPT^IVS10+16/WT^*neurons based on (**C**) immunofluorescence and (**D**) Western blotting. (**E**) Emerin transcripts are unchanged in *MAPT^IVS10+16/WT^*neurons based on ddPCR. n=3 biological replicates per group for immunofluorescence and Western blotting; n=12 biological replicates per group for ddPCR. t-test, *p≤0.05, **p≤0.01. Error bars indicate SEM.

To determine if tau-induced emerin elevation is conserved in a cellular model that does not rely on tau overexpression, we utilized iPSC-derived neurons from a patient carrying a heterozygous *MAPT IVS10+16* mutation, which causes autosomal dominant frontotemporal dementia by altering tau splicing, along with its isogenic CRISPR-Cas9-corrected control [40]. Consistent with iTau-derived neurons, we find that emerin protein is elevated in *MAPT^IVS10+16^* neurons based on immunofluorescence (**Fig. 2C**) and Western blotting (**Fig. 2D**). Emerin transcript levels are unchanged between *MAPT^IVS10+16^* neurons vs. isogenic controls based on digital droplet PCR (ddPCR) (**Fig. 2E**), suggesting a post-transcriptional mechanism underlying tau-induced elevation of emerin protein. Taken together, we find significant tau-induced elevation of emerin that is conserved between tau mutations and cellular models of tauopathy.

### Emerin regulates the actin cytoskeleton and lamin nucleoskeleton in BE(2)-C-derived neurons

While *in vitro* studies and work in epithelial cells, HeLa cells and mouse embryonic fibroblasts suggest that emerin preferentially binds to lamin A, and that disruption of B-type lamins does not alter the cellular localization or function of emerin [39,41–45], little is known regarding the function of emerin in neurons, which primarily express B-type lamins. We transfected BE(2)-C-derived neurons with GFP-tagged emerin to determine if overexpression of emerin is sufficient to induce neurotoxicity (**Supplemental Fig. 1A**). Emerin overexpression drives an increase in cytotoxicity based on increased lactate dehydrogenase (LDH), a marker of a cell damage and death, in conditioned media (**Supplemental Fig. 1B**). While LDH is elevated in conditioned media from iTau cells compared to iGFP, RNAi-mediated knockdown of emerin (emerin^KD^) (**Supplemental Fig. 1C**) does not alter secreted LDH in iGFP or iTau cells (**Supplemental Fig. 1D**), consistent with previous reports showing that emerin loss in *C. elegans* embryos, HeLa cells, mice, and mouse embryonic fibroblasts does not decrease viability [21,46–48].

We next determined if emerin regulates the actin cytoskeleton and lamin nucleoskeleton in BE(2)-C-derived neurons. Based on phalloidin staining, which labels F-actin, we find that overexpression of GFP-emerin is sufficient to increase overall levels of F-actin and F-actin thickness in neurons (**Fig. 3A**). iTau neurons also have increased levels of F-actin (**Supplemental Fig. 1E**), consistent studies in tau transgenic *Drosophila* reporting increased F-actin and the ability of tau to promote actin bundling *in vitro* [17,49,50]. Having found that induction of pathogenic tau causes emerin elevation, and that emerin elevation is sufficient to increase F-actin, we next investigated if emerin knockdown impacts F-actin in iGFP or iTau-derived neurons. While emerin knockdown does not alter total levels of F-actin, we find that the overall organization of the actin cytoskeleton is dramatically altered upon emerin knockdown in both iGFP and iTau-derived neurons (**Fig. 3B**).

**Figure 3:**
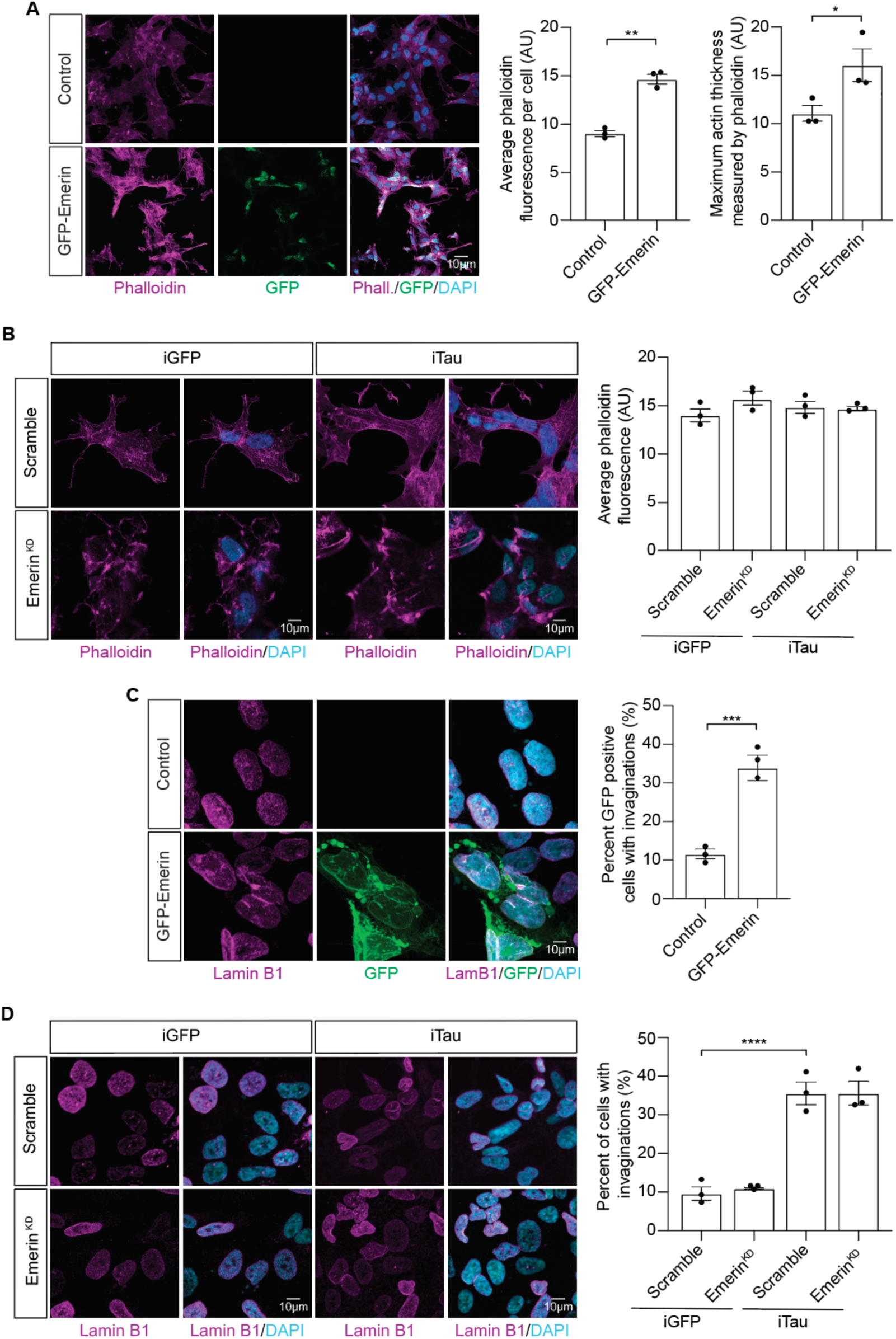
Emerin regulates the actin cytoskeleton and lamin nucleoskeleton in BE(2)-C-derived neurons. (**A**) Emerin overexpression causes an increase in F-actin levels and thickness based on phalloidin staining. (**B**) Total levels of F-actin are unchanged in iGFP and iTau cells in response to emerin knockdown, but actin structure is severely altered. (**C**) Emerin overexpression is sufficient to induce nuclear invagination based on lamin B1 immunofluorescence. (**D**) Nuclear invaginations are unchanged in iGFP and iTau cells in response to emerin knockdown. n=3 biological replicates per group. t-test or two-way ANOVA. *p≤0.05, **p≤0.01, ***p≤0.001, ****p≤0. 0001. Error bars indicate SEM.

Considering previous evidence suggesting that tau-induced overstabilization of F-actin drives nuclear invagination [15], we next determined if emerin overexpression is sufficient to induce nuclear pleomorphism using immunofluorescence-based visualization of lamin B1. We detect a robust increase in nuclear invaginations in BE(2)-C-derived neurons overexpressing emerin (**Fig. 3C**), whereas emerin knockdown does not reduce the extent of nuclear invagination in iGFP or iTau-derived neurons (**Fig. 3D**). Taken together, our findings suggest that the previously-reported role of emerin as a cytoskeletal and nucleoskeletal regulator [21,23,51–53] does indeed extend to cells that predominantly express B-type lamins.

### Emerin has increased interaction with cytoskeletal proteins in iTau-derived neurons

Given that emerin interacts with proteins along the inner nuclear membrane but translocates to the cytosol to regulate actin in response to mechanical stress, we next determined whether pathogenic tau alters emerin binding partners. Immunoprecipitation of emerin from iGFP and iTau- derived neurons (**Supplemental Fig. 2A**) followed by mass spectrometry identifies 3,800 proteins per sample, with high group correlation among replicates (**Supplemental Data 2**). Induced expression of pathogenic tau markedly alters proteins that interact with emerin (**Fig. 4A**).

**Figure 4:**
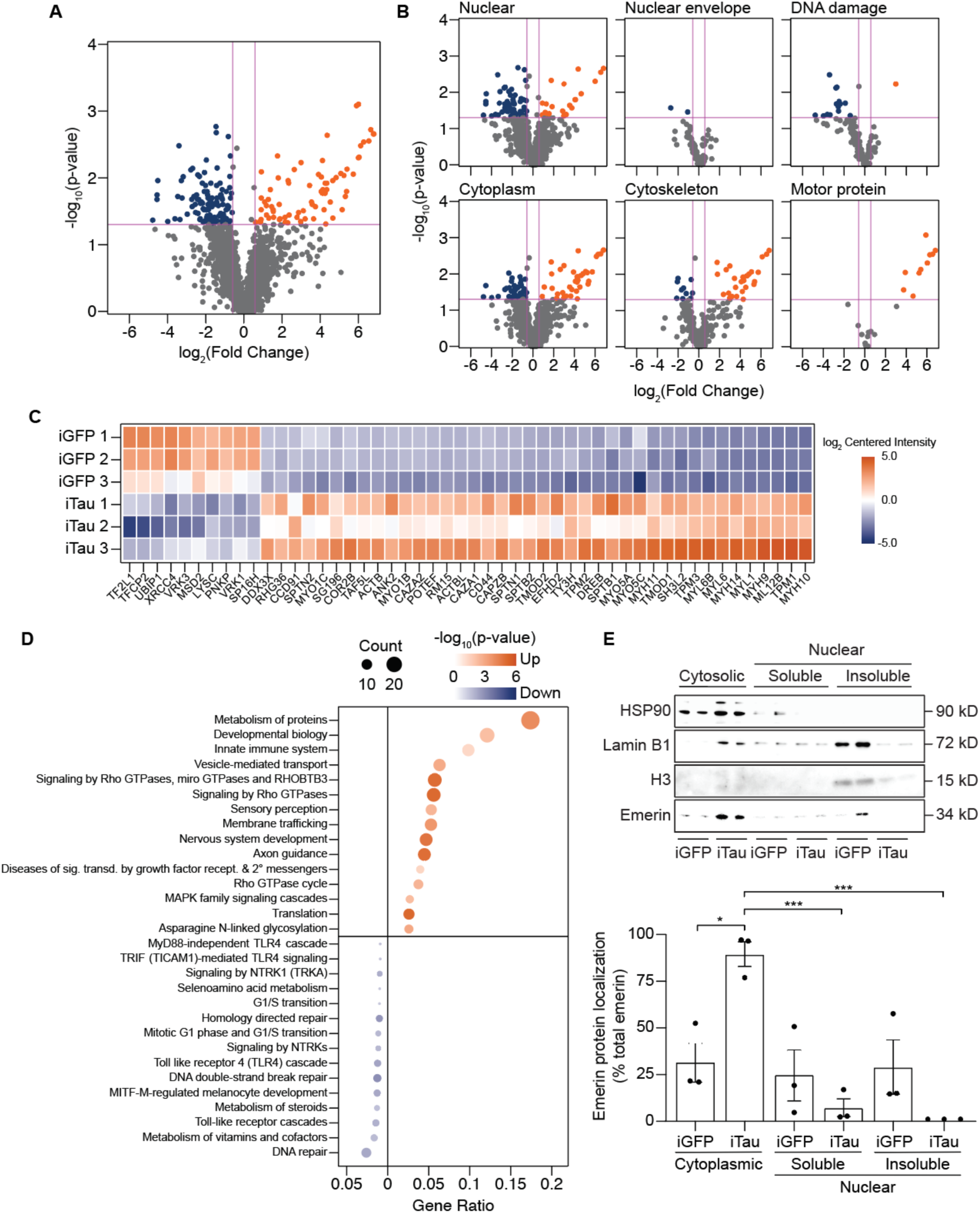
Emerin has increased interaction with cytoskeletal proteins in iTau neurons. (**A**) Volcano plot showing the altered interactome of emerin in iTau neurons compared to iGFP based on mass spectrometry. (**B**) Emerin interactome stratified by cytoplasm (GO:0005737), cytoskeleton (GO:0005856), motor proteins (GO:0003774), nuclear (GO:0005634), nuclear envelope (GO:0005635), and DNA damage (GO:0006974) GO terms. (**C**) Top 50 significant proteins with the largest change in abundance in the emerin interactome in iTau cells. (**D**) Pathway analysis of emerin-associated proteins that are significantly altered in iTau neurons. (**E**) Cytosolic emerin is elevated in iTau cells based on cellular fractionation. n=3 biological replicates per group. Log_2_FC for significance is +/- 0.585, and p≤0.05. t-test, *p≤0.05, ***p≤0.001.

Nuclear emerin regulates chromatin architecture and functions as a co-transcription factor [14,23,53]. Based on GO analysis, we find that emerin has decreased interaction with nuclear proteins, proteins at the nuclear envelope, and proteins that regulate the DNA damage response in iTau cells (**Fig. 4B**). In line with a potential relocalization of emerin from the nucleus into the cytoplasm in the context of tauopathy, we detect increased interaction between emerin and cytoplasmic, cytoskeletal, and motor proteins in iTau cells (**Fig. 4C**). Consistent with proximity-labeling experiments identifying emerin as a tau-interacting protein [32], we find increased interaction between emerin and tau in iTau-derived neurons. *In vitro* studies in HeLa cells report that outer nuclear membrane-localized emerin can bind the capping end of F-actin, which stabilizes actin filaments and facilitates a higher rate of polymerization [51]. We detect significantly higher levels of emerin interaction with actin capping proteins in iTau-derived neurons, most notably tropomodulin 1 and 2 (TMOD1, TMOD2), F-actin-capping protein subunit alpha (CAZA), and F-actin-capping protein subunit beta (CAPZB). Interaction between emerin and CAZA and CAPZB is unexpected, given previous *in vitro* evidence that emerin does not seed actin polymerization at the barbed end [51]. Induction of pathogenic tau also significantly increases interactions between emerin and various subtypes of myosin. Studies in fibroblasts, primary myoblasts, and colorectal adenocarcinoma cells suggest that myosin-emerin interactions are critical for nuclear positioning, anchoring cortical actin near the nuclear envelope, and adapting to extracellular traction forces [54–57]. Pathway analysis further reveals increased interactions with Rho GTPase signaling machinery, consistent with the greater interaction between emerin and cytoskeletal proteins (**Fig. 4D**).

We performed cellular fractionation to directly determine if tau causes emerin to accumulate in the cytosol. Consistent with mass spectrometry analysis, emerin is significantly elevated in the cytosol of iTau-derived neurons, while soluble levels of nuclear emerin are equal between iGFP and iTau (**Fig. 4E**). Insoluble nuclear emerin is undetectable in iTau-derived neurons. Adding to previous studies reporting that tau causes an overall decrease in nuclear B- type lamins [15], we detect a reduction of insoluble lamin, presumably in intermediate filament form, in iTau-derived neurons, alongside an increase in lamin B1 in the cytosol. Taken together, these analyses suggest that the intracellular mechanical environment is significantly disrupted in tauopathy and that pathological tau causes emerin to accumulate in the cytoplasm, where it regulates cytoskeletal structure.

## DISCUSSION

Extensive evidence from multiple model systems suggests that various forms of pathogenic tau disrupt the neuronal cytoskeleton and nucleoskeleton [15–17,20,27,58]. Mechanistically, studies in *Drosophila* demonstrate that pathological tau causes overstabilization of F-actin [17], which disrupts the LINC complex and destabilizes the lamin nucleoskeleton, ultimately contributing to nuclear invagination, blebbing and neuronal death [15]. Complementary work in iPSC-derived neurons reveals that tau pathology impairs the microtubule network, which also contributes to nuclear invaginations and blebbing [16]. In primary neurons, optogenetic induction of tau oligomerization deforms the nuclear envelope and promotes direct interactions between tau multimers, lamins, and nuclear pore proteins [26].

Given the well-established effects of tau on the cyto- and nucleoskeleton, alongside our previous finding that pathogenic forms of tau reduce nuclear tension [20], we were interested to find that the mechanically-sensitive protein emerin is elevated in iTau neurons and in iPSC-derived neurons carrying a *MAPT* mutation. While few studies have focused on the role of emerin in neurons, emerin was recently identified as the top activity-induced protein that is elevated in response to visual experience in mice [24]. Synthesis of emerin protein was found to occur rapidly in response to visual experience and to be independent of transcription, suggesting that an existing pool of emerin RNA is leveraged to quickly translate emerin protein in response to neuronal activity. In the current study, we also find that emerin elevation in *MAPT* mutant iPSC-derived neurons also occurs in the absence of a concomitant increase in emerin RNA, suggesting a post-transcriptional mechanism whereby tau drives elevation of emerin protein.

While we do not currently know the consequences of tau-induced emerin elevation, we find that overexpression of emerin in BE(2)-C derived neurons is sufficient to elevate F-actin and drive nuclear envelope invagination, cellular phenotypes that also occur in tauopathy [15–17,20,27,58], and is toxic to neurons based on elevated levels of LDH in conditioned media. In mouse neurons, emerin overexpression is reported to decrease protein synthesis, including proteins that promote synaptogenesis [24]. While the literature is mixed regarding the effects of tau on protein translation depending on model organism and disease stage [59–63], we speculate that tau-induced emerin elevation impacts the overall translation rate in neurons.

Given the robust effects of neuronal emerin overexpression observed in the current study, we were surprised to find that emerin knockdown disrupts actin organization in iGFP and iTau neurons but has no effect on nuclear morphology or neurotoxicity. It is possible that LEM-domain proteins LAP2 or MAN1 compensate for emerin knockdown in neurons, as has been reported in mouse muscle and liver [64]. Our findings regarding the effects of emerin knockdown on F-actin are nevertheless aligned with studies in fetal lung fibroblasts in which acute downregulation of emerin causes disorganized actin networks, often with peripheral actin bundling [23].

Studies in non-neuronal cells establish that emerin interacts with lamins, actin, and motor proteins to regulate mechanotransduction, and that emerin alters its cellular localization to respond to mechanical stress s[8,21,23,37–39]. We find that emerin is reduced in the nucleus and is significantly elevated in the cytosol in iTau cells compared to control, in line with studies in HEK293 cells reporting reduced nuclear emerin upon tau induction [65]. Based on mass spectrometry, we find that emerin has increased interactions with many cytoskeletal proteins, including several myosin subtypes, actin-binding proteins, and tau itself, in iTau neurons, and reduced interaction with nuclear proteins. Taken together, our findings support a model in which pathological forms of tau increase cytoskeletal force on the nucleus, driving emerin elevation and relocalization from the nucleus to the cytosol, where it interacts with actin, myosin and other factors to reorganize the cytoskeleton in response to nuclear force.

To date, most functional studies of emerin have been conducted in the context of non-neuronal cells. Our analyses provide new insight into the function and negative consequences of emerin overexpression in neurons and identify emerin as a candidate mediator of mechanical dysfunction in tauopathy. This work provides a strong foundation for future studies exploring how tau pathology alters the mechanical landscape in neurons and highlights emerin as a potential effector of tau-induced toxicity.

## MATERIALS AND METHODS

### Cell culture

Human neuroblastoma cells (BE(2)-C; ATCC #CRL-2268) were cultured in a 1:1 mixture of Eagle’s minimum essential medium (EMEM) and F12 medium supplemented with 10% tetracycline-free fetal bovine serum (FBS) and 1% penicillin-streptomycin. Details on the development of stable cell pools of BE(2)-C iTau (tau^R406W^) and iGFP cells are previously reported [20]. Expression of tau^R406W^ or GFP was induced with 1 µg/mL doxycycline hyclate in DMSO.

Human iPSC lines were obtained from the Tau Consortium cell line collection (https://www.neuralsci.org/tau) [40]. iPSC lines were maintained in complete mTeSR1 (#85850, Stem Cell Technology) growth medium on Corning® Matrigel® hESC-Qualified Matrix (#354277, Corning) and passaged every three to five days using ReLeSR™ (#05872, Stem Cell Technology). Cortical neurons were generated (dx.doi.org/10.17504/protocols.io.x9bfr2n) with minor modifications in the generation of neuronal aggregate formations and neuronal maturation. Differing from the protocol cited, iPSCs were plated on Nunclon Sphera 96 U bottom plates (#174929, ThermoFisher Scientific) and fed with STEMdiff™ Neural Induction Medium (NIM) + 1 µl/ml ROCK inhibitor Y-27632 (#72302, Stem Cell Technology) and plated at 3.0x10^6^ cells per well. From days two to six, fresh NIM media supplemented with dorsomorphin (#P5499, Sigma-Aldrich) and SB-431542 (#1614, Tocris) was added to cells daily. During neuronal maturation, the neurons were fed based on the Stem Cell Technology protocol: 10 ml of the BrainPhys™ Neuronal Medium (#05790, Stem Cell Technology) supplemented with 200 µl of NeuroCult^TM^ SM1 Neuronal Supplement (#05711, Stem Cell Technology), 200 µl of N2 Supplement-A (#07152, Stem Cell Technology), 20 ng/ml human recombinant brain-derived neurotrophic factor (BDNF) (#78005, Stem Cell Technology), 1 mM dibutyryl-cAMP (#73886, Stem Cell Technology) and 200 nM ascorbic acid (#72132, Stem Cell Technology). Neurons were fed every Monday, Wednesday, and Friday for eight weeks.

### iGFP and iTau neuronal differentiation

Cells were plated at 5.0x10^4^ in 6-well plates for 24 hours in a 1:1 mixture of EMEM and F12 medium supplemented with 10% tetracycline-free FBS and 1% penicillin-streptomycin. From days two to six, cells were cultured in EMEM, 1% tetracycline-free FBS, 1% penicillin-streptomycin, and 10 μm retinoic acid (#R2625, Sigma-Aldrich). Cells were fed every other day until day six, at which point 1 µg/mL doxycycline hyclate dissolved in DMSO was added to the media to induce GFP or tau^R406W^ expression.

### Mass spectrometry sample preparation

One T75 (global proteomics), or 12 T75s (emerin interactome analysis) were cultured to 95% confluency per replicate. Cells were washed with dPBS prior to harvesting, after which 0.25% trypsin was added to each flask for three minutes. 9 mL of F12 and 1 mL of FBS were added to the flask to dilute the trypsin, and cells were centrifuged for five minutes (1,000 rpm, room temperature). Cell pellets were washed with 10 mL of dPBS and centrifuged for five minutes (1,000 rpm, room temperature) to re-pellet. Cell pellets were flash frozen and stored at -80 °C.

Cell samples were randomized for preparation and DIA-MS analysis. Cells were lysed in a buffer containing 5% SDS/50 mM triethylammonium bicarbonate (TEAB) in the presence of protease and phosphatase inhibitors (#78440; ThermoFisher Scientific) and nuclease (#88700, ThermoFisher Scientific). Aliquots of the lysates containing 100 µg of protein (#R33200, ThermoFisher Scientific) were mixed with a buffer containing 10% SDS/50 mM TEAB, reduced with tris(2-carboxyethyl)phosphine hydrochloride (TCEP), and alkylated in the dark by iodoacetamide. After quenching with dithiothreitol, 12% phosphoric acid solution was added to each sample, and the mixtures were applied to S-Traps (mini; ProtiFi) for tryptic digestion (#V5111; Promega) in 50 mM TEAB. Peptides were eluted from the S-Traps sequentially with 50 mM TEAB, 0.2% formic acid, and 0.2% formic acid in 50% aqueous acetonitrile. Pooled eluates were dried by vacuum centrifugation, redissolved in the starting HPLC mobile phase (3% B, see Mass spectrometry analysis (DIA-MS) and data processing), and quantified using Pierce™ Quantitative Fluorometric Peptide Assay (#23290, ThermoFisher Scientific). Sample injections were 2 µg of peptides in 5 µL.

To extract protein lysate for emerin immunoprecipitation, cell pellets were placed on ice in 500 μL of ice-cold lysis buffer consisting of 50 mM Tris-HCl pH 7.4, 300 mM NaCl, 0.3% v/v Triton X-100, 5 mM EDTA, 1 mM DTT, 100 μM PMSF, 1 μg/mL pepstatin A and 1X Halt protease inhibitor cocktail (#78430, ThermoFisher Scientific). Once thawed, cell pellets were resuspended by vortexing. Samples were incubated on ice for ten minutes, sonicated on ice for a total processing time of 10 seconds (intervals of 0.5 seconds ON/30 seconds OFF), and centrifuged for 30 minutes (16,000xg, 4 °C) to pellet insoluble material. Each sample was quantified via Pierce BCA assay (#23225, ThermoFisher Scientific), then diluted to ∼7 mg/mL and frozen at -80 °C.

For emerin immunoprecipitation, 50 μL per sample of Dynabeads™ Protein-G (#10003D, ThermoFisher Scientific) was washed with 500 μL of PBS + 0.05% Tween 20. The tubes were then placed in a magnetic tube holder, and the wash buffer was aspirated. Washed Dynabeads were incubated at 4 °C on a rotator with 1 μg/μL primary antibody (ab40688, Abcam) in PBS for six hours. After unbound antibody was removed, 200 μL of protein lysate was added and incubated at 4 °C on a rotator overnight. Dynabeads were then washed three times in 0.05% PBS Tween 20, and protein was eluted in 15 μL of 0.5 mM glycine pH 2.8. The elution was added to 15 μL of 2x Laemmli with β-mercaptoethanol and boiled at 70 °C for ten minutes. Samples were stored at -80 °C.

Samples from the emerin co-immunoprecipitation experiment were blocked by replicate and randomized within each replicate for preparation and DIA-MS analysis. The eluates (30 µL) were mixed with 12% phosphoric acid solution and 10% SDS/50 mM TEAB, applied to S-Traps (micro; Protifi), reduced/alkylated with a mixture of 10 mM TCEP/25 mM 2-chloroacetamide in 50 mM TEAB, and then digested with trypsin. Peptide quantitative analysis was not conducted. Equal volumes of digest were used for DIA-MS analysis.

### Mass spectrometry analysis (DIA-MS) and data processing

DIA-MS was conducted on an Orbitrap Fusion Lumos mass spectrometer (Thermo Scientific). On-line HPLC separation was accomplished with an RSLC NANO HPLC system (Thermo Scientific/Dionex). Conditions for analysis of the cell lysates were: column, PicoFrit™ (New Objective; 75 μm i.d.) packed to 15 cm with C18 adsorbent (Vydac; 218MS 5 μm, 300 Å); mobile phase A, 0.5% acetic acid (HAc)/0.005% trifluoroacetic acid (TFA) in water; mobile phase B, 90% acetonitrile/0.5% HAc/0.005% TFA/9.5% water; gradient 3 to 42% B in 120 min; flow rate, 0.4 μL/min. Separation conditions for peptides from the co-IP experiments were: column, PepSep (Bruker; ReproSil C18, 15 cm x 150 µm, 1.9 µm beads); mobile phases A and B and gradient, same as above; flow rate, 800 nL/min. Peptides from cells were analyzed independently from the co-IP experiment. Separate pools were made of the digests of the cell lysates (0.4 µg/µL) and the co-IP digests (equal volumes of each). Aliquots of the pools (cells, 2 µg in 5 µL; co-IP digests, equal proportion in 5 µL) were analyzed using three stages of gas-phase fractionation (395–605 m/z, 595–805 m/z, 795–1005 m/z, staggered) and 4-m/z windows (30k resolution for precursor and product ion scans, all in the orbitrap). The resulting three data files were used to create an empirically-corrected DIA chromatogram library [66] by searching against a Prosit-generated predicted spectral library [67] based on the UniProt Human protein database [UniProt_Human_ref 9606_20220216 (20,588 sequences; 11,394,277 residues)]. MS data for the experimental samples were acquired using 8-m/z windows (400–1000 m/z; staggered; 30k resolution for precursor and product ion scans) and searched against the chromatogram library. *Scaffold DIA* (Proteome Software; cells, v3.3.1; co-IP, v3.4.1) was used for all DIA-data processing: fixed modification, cysteine carbamidomethylation; proteolytic enzyme, trypsin with one missed cleavage allowed; peptide mass tolerance, ±10.0 ppm; fragment mass tolerance, ±10.0 ppm; charge states, 2+ and 3+; peptide length, 6–30. Files were separately processed for cells and co-IP digests. Peptides identified in each sample were filtered by *Percolator* [68] to achieve a maximum FDR of 1%. Individual search results for each sample type were combined, and peptide identifications were assigned posterior error probabilities and filtered to an FDR threshold of 1% by *Percolator*. Peptide quantification was performed by *Encyclopedia* [66] based on the three to five highest quality fragment ions. Only peptides that were exclusively assigned to a protein were used for relative quantification.

### Western blotting

Cells were harvested at ∼90% confluency using RIPA with 1x Halt protease inhibitor (#78430, ThermoFisher Scientific) and incubated at 4 °C for 30 minutes with gentle rocking. Cell lysates were centrifuged for 20 minutes (12,000 rpm, 4 °C). A Pierce BCA assay (#23225, ThermoFisher Scientific) was performed on the supernatants to quantify protein concentration. Protein lysates were boiled in 2x Laemmli buffer for five minutes, centrifuged for one minute (12,000 rpm, room temperature) then loaded onto a 4–20% SDS–PAGE gel with 20 µg of protein loaded per well. Equal loading was assessed by Ponceau S staining of nitrocellulose membranes after transfer. Membranes were blocked in PBS containing 0.05% Tween 20 and 2% milk then incubated with primary antibodies overnight at 4 °C (**Supplemental Table 1**). After washing, membranes were incubated with HRP-conjugated secondary antibodies for two hours at room temperature. Blots were developed with an enhanced chemiluminescent substrate. Densitometry was performed using ImageJ.

### Immunofluorescence

Cells were plated in 12-well plates on 20 mm coverslips prior to staining. Cells were fixed in 100% ice-cold methanol at room temperature for five minutes then washed three times with 0.01% Tween 20 in PBS for five minutes per wash. After washing, cells were permeabilized in 0.01% PBS Triton X-100 with 1% bovine serum albumin (BSA) for 15 minutes at room temperature, washed three times with 0.01% Tween 20 in PBS, then blocked for 30 minutes in PBS with 1% BSA and 0.01% Tween 20. Primary antibodies were diluted in 1% BSA and incubated with cells overnight at 4 °C (**Supplemental Table 1**). The following day, cells were washed with 0.01% Tween 20 in PBS then incubated with secondary antibody for one hour at room temperature. After washing, cells were incubated with 1X DAPI for two minutes to stain nuclei then mounted onto glass slides with Vectashield (#H-1000-10, Vectorlabs). To visualize actin, cells were stained with Acti-Stain 555 Phalloidin (#PHDH1-A, Cytoskeleton) for 30 minutes prior to DAPI staining (**Supplemental Table 2**). Cells were visualized by confocal microscopy (Zeiss LSM 880 or Zeiss LSM 980). ImageJ was used for analysis.

To calculate the percentage of nuclei containing nuclear invaginations, 100 cells per replicate were scored for the presence of invaginations. Quantification of average fluorescence of emerin and phalloidin in iGFP/iTau cells was performed using ImageJ by measuring mean intensities. Quantification of emerin in *MAPT^IVS10+16^*neurons was completed by measuring mean intensities within MAP2-positive neurons using ImageJ. The ImageJ MorphoLibJ plugin was used to calculate the thickness of actin filaments.

### ddPCR

One 6-well plate was collected per biological replicate using TRIzol (#15596026, ThermoFisher Scientific) according to the manufacturer’s protocol. RNA quantities were measured using a Nanodrop8000 spectrophotometer (ThermoFisher Scientific). Equal quantities of RNA were added to a reverse transcription reaction (#4368814, ThermoFisher Scientific). Equal quantities of cDNA were loaded into QX200 Droplet PCR System (Bio-Rad). Probes for emerin and TATA-box binding protein (TBP) were predesigned by Bio-Rad.

### Emerin siRNA and GFP-emerin transfections

1% penicillin-streptomycin was removed from the media on day four of differentiation for siRNA transfection and day five for GFP-emerin transfection. Media was replaced with Gibco Opti-MEM reduced serum medium (#31985088, ThermoFisher Scientific). Cells were transfected in a 6- well plate with 100 nM of emerin Silencer^®^ Select siRNA (#4392420, ThermoFisher Scientific, Assay ID S4647 or S4645) with 9 μL/well of Lipofectamine RNAiMAX Transfection Reagent (#13778030, ThermoFisher Scientific) following the manufacturer’s protocol. For GFP-emerin transfections, cells were transfected with 6 μg of GFP-emerin (plasmid #61985, Addgene, DOI: 10.1083/jcb.201009068) with 9 μL/well of Lipofectamine 2000 (#11668019, ThermoFisher Scientific). Cells were incubated with siRNA/Lipofectamine or GFP-emerin/Lipofectamine for 24 hours and six hours, respectively. Transfection media was replaced by EMEM, 1% tetracycline-free FBS, 10 μm retinoic acid (#R2625, Sigma-Aldrich), and 1 µg/mL doxycycline hyclate in DMSO. Cells were collected for experimentation on day seven.

### Statistical analyses

For mass spectrometry, the false discovery rate (FDR) was set to 1.0%. Samples with one or more missing values, and any proteins that had a peptide count of less than two were excluded. A p≤0.05, and a Log2FC of -0.585 and 0.585 were considered significant. Correlations among and between samples were determined using the Pearson correlation coefficient. Proteins that were significantly up- and down-regulated or had a significant increase or decrease in interaction with emerin were entered into *Reactome* (Reactome.org) for pathway analysis. All plots were created using R Studio.

For immunofluorescence, samples were randomized during image acquisition and were analyzed by ImageJ. All cell replicates reflect different passage numbers from different flasks. Each experiment was completed in triplicate on different days. Western blots were analyzed by band densitometry using ImageJ. A p-value <0.05 by ANOVA and Tukey’s test for multiple comparisons and a student’s t-test for single comparisons was considered significant. GraphPad Prism was used for the analysis of immunofluorescence, Western blotting, and ddPCR experiments.

## ACKNOWLEDGEMENTS

A portion of this work was performed at Brown University with support from the Department of Molecular Biology, Cell Biology & Biochemistry. We thank Dr. Katherine Wilson for troubleshooting the emerin immunoprecipitation protocol. Mass spectrometry analyses were conducted at the University of Texas Health San Antonio (UTHSA) Institutional Mass Spectrometry Laboratory, supported in part by UTHSA and by the University of Texas System Proteomics Core Network for the purchase of the Orbitrap Fusion Lumos mass spectrometer.

## AUTHOR CONTRIBUTIONS

Conceptualization: C.S., B.F., Methodology: C.S., S.P., D.M., M.L., M.G.T., S.R.P., Validation: C.S., S.R.P., Formal analysis: C.S., M.L., E.J.S., Investigation: C.S., S.P., D.M., M.L., M.G.T., S.R.P., Resources: S.T.W., B.F., Data Curation: C.S., S.T.W., B.F., Writing – original draft preparation: C.S., Writing – review and editing: C.S., S.P., D.M., M.L., M.G.T., S.R.P., E.J.S., S.T.W., B.F., Visualization: C.S., M.L., E.J.S., Supervision: S.T.W., B.F., Project administration: B.F., Funding acquisition: C.S., E.J.S., S.T.W., B.F.

## DATA AVAILABILITY

Mass spectrometry datasets will be made publicly available through the ProteomeXchange upon acceptance of the manuscript.

## SUPPLEMENTAL FIGURES

**Supplemental Figure 1:**
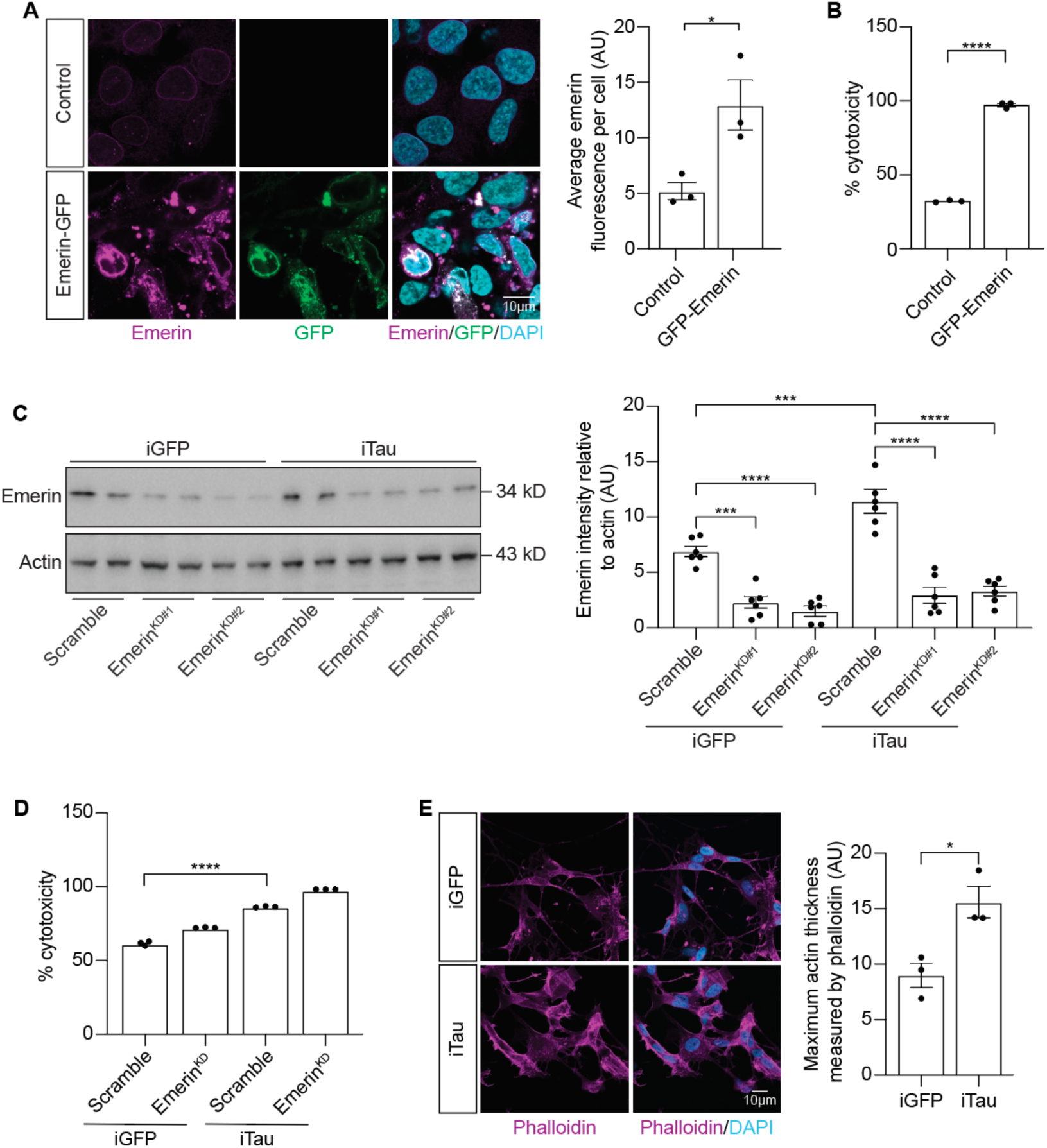
Emerin overexpression drives cytotoxicity in BE(2)-C-derived neurons. (**A**) GFP-emerin BE(2)-C-derived neurons have elevated levels of emerin based on immunofluorescence. (**B**) Significantly increased LDH in conditioned media from GFP-emerin BE(2)-C-derived neurons. (**C**) Validation of Emerin^KD^ based on Western blotting and densitometry. (**D**) LDH in conditioned media from iGFP and iTau-derived neurons is unchanged in response to emerin knockdown. (**E)** Phalloidin staining indicates that induction of pathological tau thickens the actin cytoskeleton based on fluorescence microscopy. n=3 biological replicates per group. t-test, *p≤0.05, ***p≤0.001, ****p≤0.0001. Error bars indicate SEM.

**Supplemental Figure 2:**
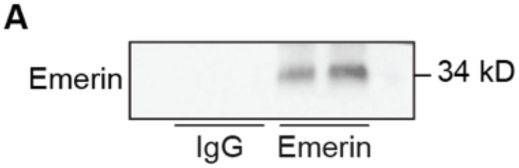
Emerin immunoprecipitation. (**A**) Coomassie stain reveals effective immunoprecipitation of emerin using emerin antibody-loaded Dynabeads™ Protein-G; IgG-loaded Dynabeads serve as a negative control.

## SUPPLEMENTAL TABLES

**Supplemental Table 1:** Antibody sources and concentrations.

| **Antibody** | **Catalog Number** | **Dilution** |
| --- | --- | --- |
| Anti-Emerin (rabbit polyclonal) | ab153718 | IHC<br>1:500/WB:1:5,000 |
| Anti-Emerin (rabbit polyclonal) | ab40688 | IP: 1 µg/µL |
| Anti-MAP2 (mouse monoclonal) | Invitrogen (13-1500) | IHC:1:200 |
| Anti-Cleaved Caspase-3 (rabbit polyclonal) | Cell Signaling Technology #9661 | IHC:1:400 |
| Anti-lamin B1 (mouse monoclonal) | ab8982 | IHC: 1:100 |
| Anti-lamin B1 (rabbit polyclonal) | ab16048 | WB:1:500 |
| Anti-HSP90 (rabbit polyclonal) | Cell Signaling Technology #4874 | WB:1:1000 |
| Anti-Actin (mouse monoclonal) | Developmental Studies Hybridoma Bank (DSHB) JLA20 | WB:1:1000 |
| Rabbit IgG Recombinant Rabbit Monoclonal Antibody | Invitrogen #PSH04-42 | IP: 1 µg/µL |
| Anti-Histone3 H3 Rabbit Polyclonal Antibody | Ab1791 | WB:1:2000 |
| Goat Anti-Mouse IgG (H+L)-HRP | SouthernBiotech #1036-05 | WB:1:20,000 |
| Goat Anti-Rabbit IgG (H+L)-HRP | SouthernBiotech #4030-05 | WB:1:20,000 |
| Goat Anti-Rabbit IgG (H+L) 488 | ThermoFisher Scientific #A-11008 | IHC:1:200 |
| Goat Anti-Rabbit IgG (H+L) 555 | ThermoFisher Scientific #A-21428 | IHC:1:200 |
| Goat Anti-Mouse IgG (H+L) 488 | ThermoFisher Scientific #A-11001 | IHC:1:200 |
| Goat Anti-Mouse IgG (H+L) 555 | ThermoFisher Scientific #A-21422 | IHC:1:200 |

**Supplemental Table 2:** Cellular dyes, sources and concentrations.

| Dye | Catalog Number | Concentration |
| --- | --- | --- |
| Acti-Stain 555 | Cytoskeleton Inc. #PHDH1-A | 100 nM/coverlip |
| DAPI | ThermoFisher Scientific #62248 | 300 nM/coverlip |

## SUPPLEMENTAL DATA

**Supplemental Data 1: DIA-MS analysis of iGFP and iTau cells.** n=3 replicates per condition.

**Supplemental Data 2: Emerin-interacting proteins in iGFP and iTau BE(2)C-derived neurons based on DIA-MS.** n=3 replicates per condition.

